# Fleas (Siphonaptera) are Cretaceous, and Evolved with Theria

**DOI:** 10.1101/014308

**Authors:** Qiyun Zhu, Michael Hastriter, Michael Whiting, Katharina Dittmar

## Abstract

Fleas (order Siphonaptera) are highly-specialized, diverse blood-feeding ectoparasites of mammals and birds with an enigmatic evolutionary history and obscure origin. We here present a molecular phylogenetic study based on a comprehensive taxon sampling of 259 flea taxa, representing 16 of the 18 extant families of this order. A Bayesian phylogenetic tree with strong nodal support was recovered, consisting of seven sequentially derived lineages with Macropsyllidae as the earliest divergence, followed by Stephanocircidae. Divergence times of flea lineages were estimated based on fossil records and host specific associations to bats (Chiroptera), suggesting that the common ancestor of extant Siphonaptera diversified during the Cretaceous. However, most of the intraordinal divergence into extant lineages took place after the K-Pg boundary. Ancestral states of host association and biogeographical distribution were reconstructed, suggesting with high likelihood that fleas originated in the southern continents (Gondwana) and migrated from South America to their extant distributions in a relatively short time frame. Theria (placental mammals and marsupials) represent the most likely ancestral host group of extant Siphonaptera, with marsupials occupying a more important role than previously assumed. Major extant flea families evolved in connection to post K-Pg diversification of Placentalia. The association of fleas with monotremes and birds is likely due to later secondary host association. These results suggest caution in casually interpreting recently discovered Mesozoic fossil “dinosaur fleas” of Northeast Asia as part of what we currently consider Siphonaptera.

## 1. Introduction

Extant fleas (Siphonaptera) are a relatively small insect order of blood-feeding ectoparasites on mammals and birds. The majority of ca. 2,575 species are adapted to rodents. Despite previous molecular analyses, their origin and the patterns of early diversification remain unresolved (Whiting et al., 2008). Recently however, the deep ancestry and evolution of fleas has been discussed in the context of new Mesozoic compression fossils from northeastern China. Based on a limited suite of morphological characters overlapping with extant fleas (e.g. winglessness and siphonate mouth structures), two research groups interpret the finds as ectoparasites, and subsequently place their evolutionary relations close to or within the extant Siphonaptera (Fig. 1). The presumed flea finds (also referred to as pre-fleas) are dated from the Early Cretaceous to the Mid Jurassic, thus extending the possible origins of the flea lineage significantly into the past (Gao et al., 2013; Gao et al., 2012; Huang et al., 2013a; Huang et al., 2012).

Previous to these finds, fleas were thought to have existed already in the Paleogene, based on fossils from Dominican and Baltic Amber (Beaucournu, 2003; Beaucournu and Wunderlich, 2001; Dampf, 1911; Peus, 1968). The amber specimens show little morphological difference from modern fleas, suggesting that the flea stem lineage likely began much earlier. Typically for ectoparasites, the fossil record is sparse, and no confirmed representatives of modern fleas older than the Late Eocene have been found to date (*Palaeopsylla* spp. in Baltic Amber). Other Jurassic and Cretaceous fossils with presumed siphonate mouth parts, such as *Tarwinia* (Jell and Duncan, 1986; Riek, 1970), *Strashila* (Rasnitsyn, 1992) and *Saurophthirus* (Ponomarenko, 1986) had been previously discussed as ectoparasites, and pre-fleas, but this remains controversial. In fact, the position of *Strashila* has recently been revised, placing them as aquatic or amphibious relatives to the extant dipteran Nymphomyiidae (Huang et al., 2013b). However, due to their understanding of *Tarwinia* as an ectoparasite (and a flea), Grimaldi and Engel (2005) suggested an Early Cretaceous origin of stem siphonapteroid insects.

**Figure 1.**
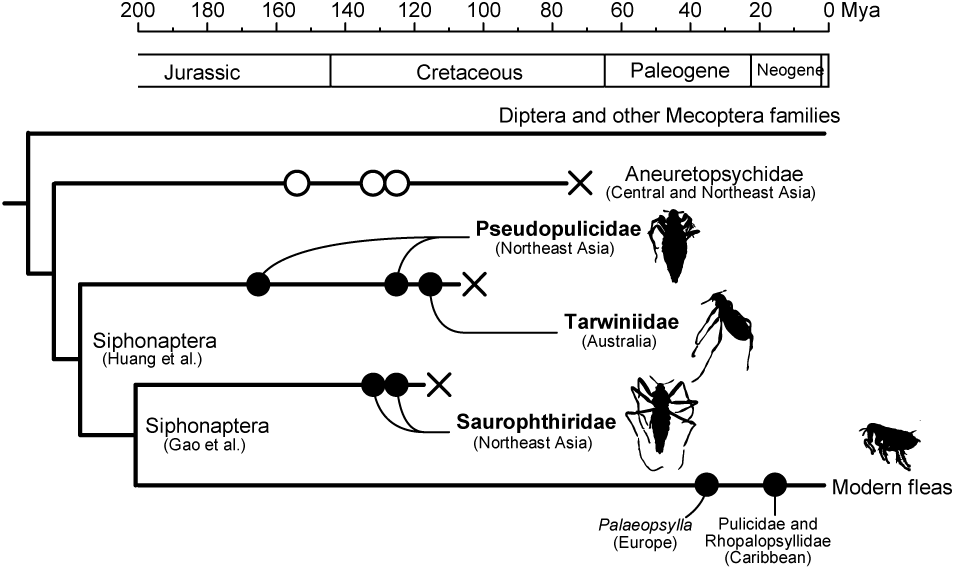
Illustration of the proposed “dinosaur flea” phylogeny. The hypothetical phylogenetic relationships among modern fleas, Mesozoic “pre-fleas” and their extant sister groups are illustrated based on Gao et al. (2013) and Huang et al. (2012, 2013a). Solid dots represent fossil records of confirmed true (extant) Siphonaptera or proposed “pre-fleas” and open dots represent those of mecopteran Aneuretopsychidae. The estimated ages of fossils are indicated by the scale bar. Locations of fossil discoveries are labeled. The three proposed extinct siphonapteran families, Pseudopulicidae (including genera *Pseudopulex*, *Hadropsylla* and *Tyrannopsylla*), Saurophthiridae (*Saurophthirus*) and Tarwiniidae (*Tarwinia*) are indicated in bold font. The silhouettes of representative specimens, as well as that of a modern flea, are illustrated beside the taxonomic names.

Phylogenetic evidence points to fleas being clearly monophyletic, and some studies have proposed a sister group relationship to the extant Boreidae (snow scorpionflies), with which they share important morphological characters (Sikes and Stockbridge, 2013; Whiting, 2002; Whiting et al., 2008). Based on the oldest known boreid fossils, *Palaeoboreus spp.* (150-140 Mya) (Sukatsheva and Rasnitsyn, 1992), the Boreidae are thought to be at least Late Jurassic in origin, suggesting the possibility that the flea stem lineage is equally old. Recent molecular studies however have suggested that Siphonaptera and Mecoptera are sister groups (McKenna and Farrell, 2010; Misof et al., 2014; Peters et al., 2014; Wiegmann et al., 2009), Divergence time analysis on holometabolous insects with limited taxonomic representation of Siphonaptera placed the split of Mecoptera and Siphonaptera into the Triassic (Wiegmann et al., 2009) or the Jurassic (Misof et al., 2014), well within the age range of the recently proposed fossil “fleas”.

The recent fossil finds of presumed Siphonaptera seemingly support a potential Jurassic origin, thus not only challenging some of the current ideas about the evolutionary timing of flea evolution, but also prior hypotheses about their ancestral host associations. Because the Mesozoic fleas co-occur in the same general fossil beds as feathered dinosaur fossils, ancestral evolution with feathered or “hairy” dinosaurs or pterosaurs has been suggested (Gao et al., 2013; Gao et al., 2012; Huang et al., 2013a; Huang et al., 2012). This conflicts with the currently accepted idea of an early siphonapteran association with mammals, which had been proposed after conducting general host-association mapping on a topology with a subset of flea lineages (Whiting et al., 2008). Specifically, proponents of the “dinosaur flea” hypothesis suggest an initial association with dinosaurs in a siphonapteran stem lineage, to be followed by a later switch to mammals (Huang et al., 2012).

By extension, this also conflicts with the idea that the ectoparasitic lifestyle of extant fleas evolved directly from free-living ancestry, as suggested by Whiting (2002). This is reflected by the novel placements of the fossils into hypothetical topological contexts of a subset of Mecoptera (Fig. 1). Huang et al. (2013a) described three genera of Pseudopulicidae fam. nov. (*Pseudopulex*, *Hadropsylla*, and *Tyrannopsylla*), which are all placed within the Siphonaptera, together with *Tarwinia* and *Saurophthirus* spp. (Saurophthiridae) (Fig. 1). On the other hand, Gao et al. (2013) place only Saurophthiridae into the Siphonaptera, and also suggest it was a transitional ancestral form of extant fleas (Fig. 1). Both research groups place the extinct Aneuretopsychidae (Ren et al., 2011) as sister group of the fleas and/or pre-fleas, similar to the results of an extensive morphological analysis of extant and extinct mecopteran lineages by Ren et al. (2009).

In light of these new suggestions, we explore an extended molecular dataset of fleas to reconstruct an updated phylogeny. For the first time, we estimate divergence times of Siphonaptera, and reconstruct ancestral host affiliation and biogeographic history.

## 2. Materials and methods

### 2.1 Taxon and Sequence Sampling

259 flea specimens representing 205 species were obtained from multiple sources, collected over two decades (1989 - 2014) (Table S1). The current dataset represents 16 of 18 flea families (88.9%, as per (Medvedev, 1994, 1998)). Voucher exoskeletons of DNA-extracted fleas were preserved, and curated as slide mounted specimens in the Monte L. Bean Life Science Museum in the Insect Genomics Collection at Brigham Young University. Eight samples from Boreidae (snow scorpionflies) were used as mecopteran outgroup (Whiting et al., 2008). DNA sequence datasets (8549 bp) of nine mitochondrial and nuclear protein-coding and ribosomal genes were prepared, containing the following genes: 12S, 16S, 18S and 28S rRNAs, elongation factor 1-alpha, histone H3, cytochrome b, cytochrome oxidase subunit I and II (Table S2). Sequence data stem from an increased gene sampling of previously published data (Whiting et al., 2008), as well as an extended species sampling (GenBank accession numbers KM890312-KM891539) (Table S1).

### 2.2 Sequence alignment

Each gene was aligned separately, using MAFFT 6.864b (Katoh, 2002; Katoh and Toh, 2008). Protein-coding genes (cytochrome oxidase subunit I and II, cytochrome b, elongation factor 1-alpha, and histone H3) were aligned with the L-Ins-i algorithm, whereas ribosomal genes (12S, 16S, 18S, and 28S rRNAs) were aligned with the Q-Ins-i algorithm. The rRNA gene alignments were further optimized with RNAsalsa 0.8.1 (Stocsits et al., 2009), which takes prior knowledge of rRNA secondary structure into account. Each alignment was filtered using the GUIDANCE algorithm (Penn et al., 2010b) implemented in the GUIDANCE server (Penn et al., 2010a) to remove ambiguously aligned regions. The insertion/deletions (indels) in the sequences were coded as binary data with SeqState 1.4.1 (Müller, 2005), using the SIC algorithm (Simmons and Ochoterena, 2000).

### 2.3 Phylogenetic Analyses

Analyses were performed on the concatenated alignment using a Bayesian approach. The alignment was partitioned by genes and codon positions. PartitionFinder was employed to compute the best partitioning scheme, as well as the best nucleotide substitution models for each partition (Lanfear et al., 2012). Bayesian information criterion (BIC) (Schwarz, 1978) was used to select the best models (Table S3).

Bayesian inference (BI) analysis was performed using Markov Chain Monte Carlo (MCMC) tree searches in BEAST 1.8.0 (Drummond and Rambaut, 2007a; Drummond et al., 2012). The relaxed clock model with uncorrelated lognormal distribution was set for the molecular clock, and the Yule process (Gernhard, 2008) was used for the tree prior. A series of test runs were performed for model optimization purpose. The posterior distributions of parameters were assessed using Tracer 1.6 (Drummond and Rambaut, 2007b). Models whose parameters consistently received low ESSs were replaced by alter native models for new analyses, to account for possible overparameterization (Table S3). With the optimized models, three independent runs with 50 million generations each were performed. Consensus trees were extracted in TreeAnnotator following the 50% majority rule, with proper burn-ins determined in Tracer 1.6. Trees generated from different runs were compared in FigTree 1.4.0 (Rambaut, 2013) to assess the consistency of topology. The consensus tree that receives the highest average posterior probability among the three runs is presented (Fig. S1). Its topology is identical to the consensus tree combined from all three runs as computed in LogCombiner (Drummond et al., 2012).

**Figure 2.**
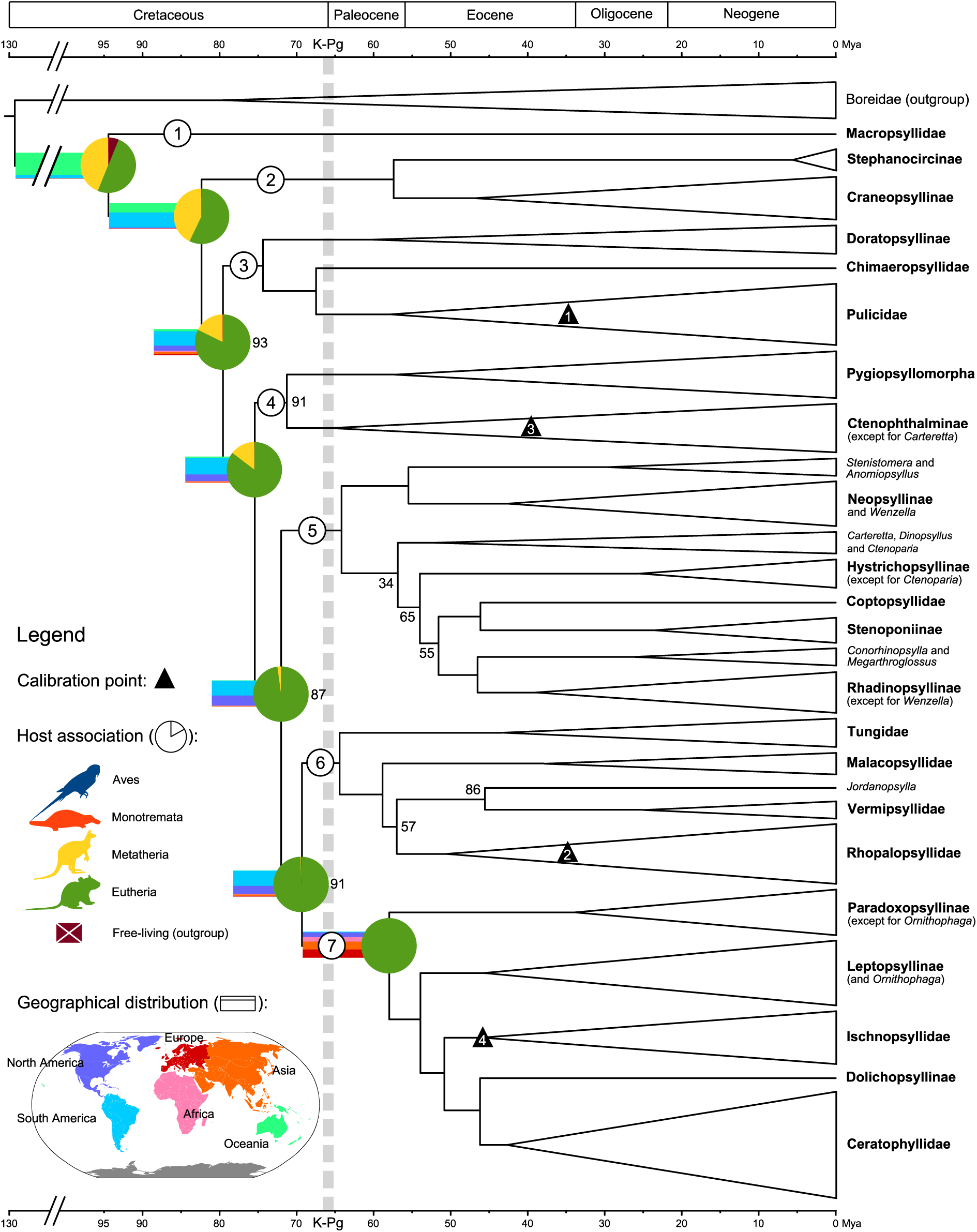
Reconstructed scenario of flea evolution: phylogeny, node ages, host associations and geographical distributions. Reconstructed Siphonaptera phylogeny is represented as a Bayesian majority consensus tree. Bayesian posterior probabilities are percentage converted, and if lower than 95 are labeled at corresponding nodes. Branch lengths are drawn to reflect the result of Bayesian divergence time estimation. The scale bars represent ages (unit: million years ago [Mya]). The indices of the seven major lineages (see 3.1) are indicated in circles. Large clades are collapsed into triangles. The height of a triangle is proportional to the square root of the number of taxa included in this clade. Higher taxonomic ranks (subfamily or above) are indicated in bold font. Reconstructed results obtained from BBM analysis are indicated at the seven major nodes. Circles (pie charts) on nodes represent estimated probabilities of occurrence of major host categories (not normalized). Bars overlapping branches represent estimated possibilities of geographical distribution by continent. For short branches, bars are extended to the left to enhance visibility. General topological locations of fossil and host-association calibrations used in the divergence time estimation analysis are represented by solid triangles, with numbers corresponding to manuscript text.

### 2.4. Divergence time estimation

The ages of origin and basal diversification of major flea groups were estimated in a Bayesian framework in BEAST. The same parameters and steps as in the standard phylogenetic analysis (see above) were used. Multiple calibrations were set up based on fossil records and knowledge of host association, following generally accepted guidelines (Benton et al., 2009; Ho and Phillips, 2009; Parham et al., 2012; Warnock et al., 2012). We chose to use fossil calibrations of confirmed true Siphonaptera only (amber specimens, see Introduction), as they impart verifiable knowledge of the existence of a particular lineage in time. Latest possible fossil ages were implemented as hard lower bounds of lognormal distributions of node age priors (Ho and Phillips, 2009; Yang and Rannala, 2006), with their 95% soft upper bounds set based on the knowledge of host association from previous records and from this study. The maximum ages of mammalian and avian hosts were uniformly referred to the values recommended by Benton and Donoghue (2007), which are based on a comprehensive review of fossil and biostratigraphic evidence. Because of the widely recognized sensitivity of divergence time estimation to the definition of priors (Nowak et al., 2013; Sauquet et al., 2012), we also tested normal and uniform distributions of node age priors for comparison to the lognormal approach (Table 1).

The following fossil calibration points were implemented:

1. Age of *Pulex*: A female flea *Pulex larimerius* was found in Dominican amber (Lewis and Grimaldi, 1997). Morphologically, the fossil is congruent with the extant genus *Pulex*, and five strands of mammalian hair provide ecological context. Although this flea fossil was placed into the Miocene, the age range of Dominican amber is controversial, with estimates running from 20-15 Mya (Iturralde-Vinent and MacPhee, 1996) to 45-30 Mya (Schlee, 1990). In order to reflect this uncertainty, the lower bound was set to be 15 Mya, and the mode (peak of highest possibility in a lognormal distribution) to be 35 Mya. Host association patterns were employed to define upper bounds. Pulicidae has a diverse and promiscuous host spectrum, with the preferred host group as rodents, followed by carnivores. Together they comprise 73% of all mammalian host records for *Pulex*. Based on Manter’s rule, stating that parasite speciation tracks or slightly lags that of hosts (Manter, 1955) it is reasonable to assume that Pulicidae (and *Pulex*) diversified with or after the diversification of either of these two mammalian groups. The maximum constraint ages of basal diversification provided by Benton and Donoghue (2007) are 65.8 Mya for Rodentia and 63.8 Mya for Carnivora, and the slightly older Rodentia was used as a conservative placement of the upper bound for the origin of *Pulex*. Recent divergence time estimates of basal diversification of Rodentia (Bininda-Emonds et al., 2007; dos Reis et al., 2012; Meredith et al., 2011; O’Leary et al., 2013) support the age range provided by Benton and Donoghue (2007). Because of the wide host range of Pulicidae, and the derived position of *Pulex* (Fig. 2), we cannot rule out the possibility of an earlier divergence on hosts other than rodents or carnivores. Therefore a comparative calibration was implemented on this node, using the maximum age of the basal diversification of placental mammals (Eutheria, 113 Mya) as an upper bound.

2. Age of *Rhopalopsyllus*: Rhopalopsyllid fossils were recorded from Dominican amber, one of which has been clearly identified as a member of the extant genus *Rhopalopsyllus* (Poinar, 1992; Poinar, 1995). Rodents are the most prevalent hosts of extant Rhopalopsyllidae (also of *Rhopalopsyllus*), although they may also occur on birds. Based on morphological characters, Poinar (1992) hypothesized that some of the fossil rhopalopsyllids were ancestrally associated with birds, similar to the extant rhopalopsyllid genus *Parapsyllus*. However, previous studies predicted rodents as the dominant ancestral host of the subclade containing *Rhopalopsyllus* spp. (Whiting et al., 2008). Therefore, the same calibration parameters were used for *Rhopalopsyllus* and *Pulex*.

3. Age of *Palaeopsylla*: Several fossil representatives of *Palaeopsylla* were found in Baltic amber (Beaucournu, 2003; Beaucournu and Wunderlich, 2001; Dampf, 1911; Peus, 1968). Despite its name, which alludes to the idea that palaeopsyllid fleas appear to be morphologically primitive, the genus does not occupy a basal position on the flea phylogeny (Whiting et al., 2008). The current taxonomic placement in the Ctenophthalmidae is debatable, since this family is not supported as a natural group (Whiting et al. (2008) and this study). The age range of Baltic amber is consistently estimated to be 35-40 Mya (Dunlop and Giribet, 2003; Ritzkowski, 1997; Weitschat and Wichard, 2010). Therefore, the lower bound was set to be 35 Mya, with the mode at 45 Mya. *Palaeopsylla* and its sister genera are predominantly associated with Soricomorpha (shrews and moles) and Rodentia. Which of the two mammalian groups is the primary host group of *Palaeopsylla* remains uncertain. Therefore we chose to use the split between Soricomorpha (Laurasiatheria) and Rodentia (Euarchontoglires) as the upper bound, which is recommended to be 113 Mya (Benton and Donoghue, 2007).

In addition to fossil records of true fleas, the fourth calibration point was based on the unique host association pattern of Ischnopsyllidae (bat fleas):

4. Age of Ischnopsyllidae: Unlike most fleas, Ischnopsyllidae are host-specific, and exclusively parasitize bats (Chiroptera) (Jameson, 1985; Krasnov, 2008). Molecular analyses clearly support the monophyly of the family (Whiting et al., 2008). Based on Manter’s first rule (Manter, 1955), we assume that the origin of Ischnopsyllidae should not be before the origin of Chiroptera, which, according to Benton and Donoghue (2007) is suggested to be not earlier than 71.2 Mya. Recent divergence time estimations including bats are generally consistent with this stem age for bats (Bininda-Emonds et al., 2007; dos Reis et al., 2012; Meredith et al., 2011; O’Leary et al., 2013). No age-indicative bat flea fossils exist, therefore, for one calibration, no minimum bound was set for this node, and a normal distribution of priors from 0 (present) to 71.2 Mya as the 95% credible interval was set.

**Table 1.**
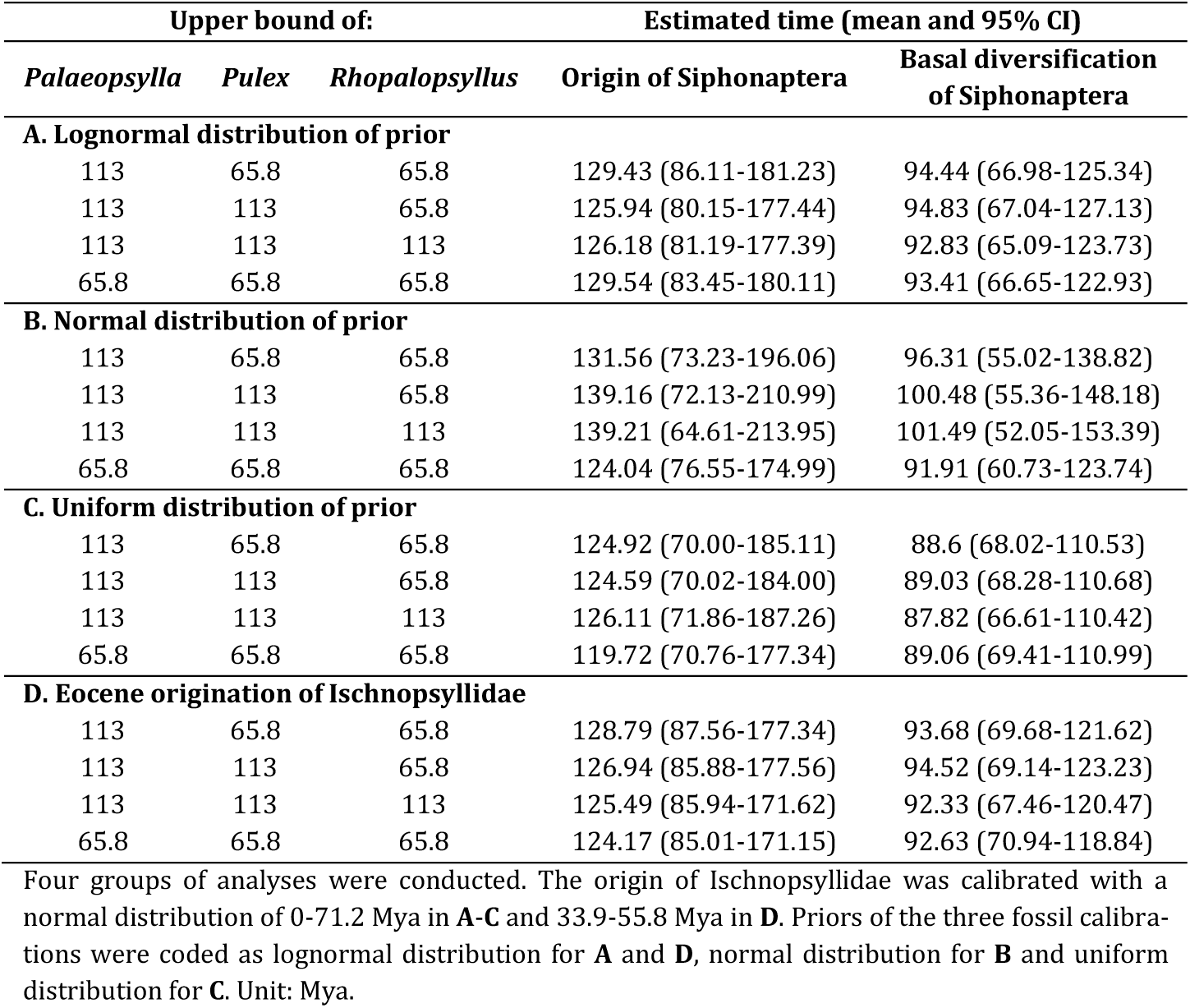
Estimated divergence times of Siphonaptera

However, despite the absence of bat flea fossils, the range of the origin of Ischnopsyllidae may be further narrowed down. Several studies suggest that bat fleas and bats coevolve to some degrees (Medvedev, 1990; Morrone and Gutiérrez, 2005; Seneviratne et al., 2009). Extant Ischnopsyllidae members have been recorded from several Chiroptera superfamilies (e.g., Molossidae and Vespertilionidae), which are proposed to have emerged from the rapid diversification of Chiroptera during Eocene (Eick et al., 2005; Simmons and Ochoterena, 2000; Simmons et al., 2008; Teeling et al., 2005). Therefore it has been suggested that Ischnopsyllidae originated during the heyday of bat speciation, that is, as early as the Eocene (Jameson, 1985; Medvedev, 1990). With this hypothesis in mind, a lognormal prior with a lower bound of 33.9 Mya, a mode of 45 Mya (Mid Eocene), and an upper bound of 71.2 Mya was used to calibrate this node for comparison.

### 2.5. Reconstruction of ancestral host association and biogeography

The host association and geographical distribution records of flea taxa were extracted from The PARHOST World Database of Fleas (www.zin.ru/animalia/Siphonaptera/index.htm), as well as Global Species (www.globalspecies.org), which is based on PARHOST. The PARHOST database is a highly curated primary resource, which has been assembled by Sergei Medvedev, under the auspices of the Zoological Institute in St. Petersburg, Russia. Pertinent information on recently described species that are not recorded in the database was manually extracted from the pertinent literature (Hastriter and Dick, 2009; Hastriter, 2009; Hastriter et al., 2009).

For the purpose of exploring patterns of host associations of the ancestral lineages of extant fleas, four general summarized host categories were considered, reflecting the breadth of known hosts: Aves, Monotremata, Metatheria and Eutheria. The three latter taxonomic units correspond to recent revisions in the mammalian tree of life as per dos Reis et al. (2012). Specifically, Eutheria is considered as comprised of placental mammals (Placentalia) plus several non-placental fossils, which group in the Eutheria (O’Leary et al., 2013). Metatheria describes the lineage leading to the extant Marsupialia.

Host association matrices were coded in continuous and categorical character states, in preparation of the analytical approaches taken (see below): (A) In the continuous matrix, for each extant flea genus represented in this study the proportion of each host category as defined by Whiting et al. (2008) was calculated from all host records as a numerical value from 0 to 1. (B) The categorical matrix was informed by the continuous data, and only hosts that were present in at least 10% or more of all host records were considered as a bona fide host category. In addition to the above described approach, two comparative matrices were generated to explore the potential influence of host species richness on ancestral host optimization. Variation in extant host species richness has been explained by differences in lineage specific rates of speciation and extinction (Mullen et al., 2011). In the context of extant flea-host associations, this process limits the number of observable ecological relationships if few host lineages survived through evolutionary time. For example, Monotremata have only 5 contemporaneous species (out of at least 12 fossil species (Musser, 2003)). One paradigm of parasite evolution is that descendant host species inherit the parasites of their ancestors. From that, an argument could be made that extant host associations should be weighted by normalizing the number of flea host species of a category against all extant species in that category. This may remedy a familiar issue of ancestral state optimization, when less speciose groups are biased against. Thus, the matrices differ from the original approach in Whiting et al. (2008) by one additional step: the number of host species per host category per flea taxon has been divided by the total number of recorded species of that category, as referring to the 2013 annual checklist of the Species 2000 database (www.catalogueoflife.org). The normalized state values were then used for the calculation of ancestral hosts.

The geographical distribution records of flea taxa were classified 1) by ecozone, and 2) by continent and coded as categorical data. (A) The ecozone coding followed the system used by Medvedev (2006), taking into account large biogeographic areas separated by significant extant migration barriers for terrestrial organisms: Palearctic, Nearctic, Afrotropic, Neotropic, Australasia and Indomalaya (Oriental). (B) The continent coding considered: Europe, Asia, Africa, North America, South America and Oceania. This system is intended to serve as an estimator of the correlation between the early evolution of fleas and plate tectonics. Occurrence of extant fleas in Antarctica was ignored, because of their sporadic, and likely secondary nature. To minimize sampling bias, each terminal (= flea species) was coded with the information of its genus.

Based on these matrices, the ancestral states of host association and biogeographic origin were reconstructed using two statistical methods: (A) The maximum parsimony (MP) approach as implemented in Mesquite 2.75 (Maddison, 1991; Maddison and Maddison, 2011) was used on continuous data (under the squared change assumption) and categorical data (under the unordered states assumption). The posterior topology from the Bayesian approach was supplied. (B) For categorical data the recently developed Bayesian Binary MCMC (BBM) approach was employed (Yu et al., 2011), as implemented in RASP (Yu et al., 2013). BBM averages the frequencies of ancestral range per node over all trees, and reports probabilities of states at ancestral nodes, rather than suggesting one ancestral state as in MP. In BBM, the F81+G model was employed to describe the character evolution. All other parameters were left at default. Two runs containing 10 MCMC chains each with a length of 100,000 generations were executed, which were sufficient for full convergence (standard deviation of split frequencies < 0.01). The first 10% samples were discarded as burn-in. The probabilities of occurrence of each host category at major nodes as computed by BBM were reported (Table S4).

## 3 Results and discussion

### 3.1. Major patterns of diversification

For the purpose of this paper, only general evolutionary insights related to the diversification of major lineages within extant Siphonaptera will be discussed. The phylogenetic analyses on an extended dataset in a Bayesian framework recover a stable topology with seven major lineages (1-7, Figs. 2 and S1), supporting once again the monophyly of Siphonaptera (Fig. 2). Nodal support measures (posterior probabilities) indicate robust values at most of the early diversifications at the infraorder level and beyond, which is a significant improvement from previous work (Whiting et al., 2008). However, some backbone nodes do not receive statistically relevant support values (posterior probabilities < 95%), making interpretations of phylogenetic relationships between lineages 2 and 3, as well as lineages 4, 5, 6, and 7 difficult. However, consistent with previous work, Pygiopsyllomorpha (part of lineage 4) and Ceratophyllomorpha (lineage 7, Fig. 2) are recovered as monophyletic, whereas Pulicomorpha and Hystrichopsyllomorpha are paraphyletic. Most importantly, there is strong support for the first divergence event as the split between Macropsyllidae (represented by *Macropsylla novaehollandiae*, lineage 1, Fig. 2) and the remainder of the fleas. This placement is incongruent with Whiting et al. (2008), which suggested that Tungidae was the first diversification event within Siphonaptera. In contrast, Tungidae is placed more apically in the tree and groups with Malacopsyllidae, Vermipsyllidae and Rhopalopsyllidae (lineage 6), all of which are families within the Pulicomorpha (*sensu* Medvedev). This different placement may result from an improved taxon sampling of this analysis, reflected in the addition of the previously not represented Malacopsyllidae and Hectopsyllini, the other tribe of the Tungidae. Similar to Whiting et al. (2008), the placement contradicts previous hypotheses of Tungidae as a sister group to Pulicidae (Hopkins and Rothschild, 1953; Lewis, 1998; Medvedev, 1998; Smit, 1988).

Lineage 2 (Fig. 2) represents the well-sampled Stephanocircidae (helmet fleas) (7/8 genera), validating previous hypotheses regarding the monophyly of the family and both subfamilies. This position suggests a closer relationship to Macropsyllidae, and an even more ancestral divergence of the family than suggested by Whiting et al. (2008). Although macropsyllid fleas share many morphological characters with Hystrichopsyllidae (Hastriter and Whiting, 2003), previous researchers have pointed to the common presence of the occipital tuber with Stephanocircidae (Hopkins and Rothschild, 1956).

Lineage 3 unites members of the Hystrichopsyllomorpha and Pulicomorpha, specifically, the monophyletic Pulicidae, the Chimaeropsyllidae (represented by one species), and the Doratopsyllinae. Similar to Whiting et al. (2008), the Chimaeropsyllidae + Pulicidae relationship has been recovered with strong support, and certain morphological characters have been discussed in the context of potential synapomorphies supporting this relationship. However, the addition of Doratopsyllinae to this lineage is new, but also strongly supported. The subfamily Doratopsyllinae is placed in the Ctenophthalmidae, underlining the urgent need for a revision of the current taxonomic arrangements in this family. Specifically, members of the Ctenophthalmidae appear scattered in 5 positions over the topology (lineages 3, 4, 5, and 6).

### 3.2. Cretaceous origin and diversification of fleas

The divergence time estimates under different distributions using the set of maximum and minimum constraints as described above (section 2d), are generally congruent and robust, and do not vary more than 7 My in either direction (mean, Table 1). Additionally, alternative parameterization introduced by both calibrations of point 4 (age of Ischnopsyllidae, bat fleas) did not seem to have a notable effect (Table 1). Results based on the parameter set as described in the first row of Table 1A are presented in Fig. 2.

In the time-calibrated Bayesian tree (Figs. 2 and S2) all stem Siphonaptera are estimated to have diversified throughout the Cretaceous, before the K-Pg extinction event (65.8 Mya). The earliest siphonapteran divergence is that of the Macropsyllidae, averaged at 95-90 Mya (Late Cretaceous), possibly starting already in the Early Cretaceous (95% CI – lower tail, Table 1). This is substantially older than the age of confirmed fossil fleas (e.g., amber fossils of Pulicidae and Rhopalopsyllidae), and indicates that extant Siphonaptera are successful survivors of a major ecological transition, which ended the reign of non-avian dinosaurs. However, we do not find support for a previously proposed Jurassic beginning of extant Siphonaptera.

It is interesting to note that except for Macropsyllidae, all other six stem lineages evolved in a narrow timeframe of 83-68 Mya, indicating a Cretaceous fuse for early intraordinal divergences, before launching into the major diversifications in the surviving stem lineages after the K-Pg boundary. Given the overall age of fleas, this may be indicative of a rapid radiation, which may be the consequence of new ecological niches (i.e. hosts) becoming available (Hayakawa et al., 2008; Price, 1980). By the Eocene (56-33.9 Mya) all extant flea families were present, and a majority had already diverged into the currently observed genus diversity.

### 3.3. Early association with Theria

Results of the ancestral host reconstruction are reported as probabilities from the BBM analysis (categorical states) and proportions from the MP analysis in Table S4. Similar optimization results were recovered at ancestral nodes by both methods. Comparing the results based on the normalized matrices to those from the original matrices, the BBM probabilities of Monotremata (which has only five extant species) are significantly elevated, while those of the other three categories do not exhibit a constant increase/decrease trend in response to the normalization. Given the scope of the paper, suggested ancestral hosts from the BBM analysis using the original host matrices were mapped onto lineage divergences in the tree (Fig. 2).

Given that fleas are ectoparasites, it is important to consider host association in the context of the timing of host radiations. Similar to previous analyses and hypotheses, the reconstructed ancestral host association at the very base of extant Siphonaptera is ambiguous respective to the host categories used, but suggests an association with mammals (Metatheria + Eutheria + Monotremata) (Krasnov, 2008; Whiting et al., 2008). Specifically, both Eutheria and Metatheria (= Theria) receive high probability (0.82 and 0.72), whereas Monotremata although present as a possible state, shows a marginal probability (4.4×10^−4^). Even when host scores are normalized to balance the current sparseness of Monotremata species, the probability (1.4×10^−3^) does not improve over that of Eutheria (0.76) and Metatheria (0.80) (Fig. 2, Table S4). This clarifies previous ideas about ancestral flea hosts. Monotremata (Prototheria) are understood as the surviving representatives of the earliest Mammalia (O’Leary et al., 2013). They are parasitized by several secondarily associated cosmopolitan pulicid fleas, as well as members of the Pygiopsyllomorpha and Stephanocircidae. The two latter groups are considered primary fleas of Monotremata, and both occupy relatively deep positions in the flea tree (Fig. 2). Considering the strong Australian connection of monotreme diversification (Archer et al., 1985; Musser, 2003), one may reasonably argue that the monotreme-flea association may be ancient, too (Grimaldi and Engel, 2005). Yet the low likelihoods of Monotremata at the basal node make this scenario less likely, which is biologically consistent with the Cretaceous age estimation of the earliest flea node being younger than the estimated Middle Jurassic split of Monotremata and Theria (Bininda-Emonds et al., 2007; dos Reis et al., 2012; Meredith et al., 2011; O’Leary et al., 2013).

As lineage diversification progressed (lineages 2-7), Eutheria became the most likely of the four flea host states considered, and any flea divergence beyond the K-Pg boundary has been mapped to this state, with significant probabilities. In other words, the ancestral state optimizations in this study lend strong support to a post-K-Pg diversification of crown fleas with Eutheria, thus coinciding with similar hypotheses regarding the crown Placentalia divergence (Bininda-Emonds et al., 2007; dos Reis et al., 2012; Meredith et al., 2011; O’Leary et al., 2013).

Birds, though being frequently parasitized by extant fleas, are not estimated to have any traceable association with early fleas (Fig. 2). In fact, all ancestral nodes associated with Aves cluster in the crown clades (not shown), and are the results of multiple independent shifts from placental mammals. This inference is consistent with previous observations and analyses (Holland, 1964; Hopkins, 1957; Krasnov, 2008; Whiting et al., 2008).

### 3.4. Biogeography supports a Gondwanan origin

Results from the reconstructed biogeographic distributions using BBM or MP methods are generally congruent. Comparatively, BBM results of continents and those of their corresponding ecozones (e.g., Oceania and Australasia) are similar, whereas this similarity is less significant in the MP results (Table S4). BBM results by continent were displayed at basal nodes of Fig. 2.

The ancestral states of stem Siphonaptera nodes are reconstructed to be Oceania and South America, with high to moderate likelihood, respectively (Fig. 2). Support for all other continents/ecozones on these early nodes is negligible. From the view point of plate tectonics, and considering the node ages from the divergence time estimation, Australia and South America together translate into part of the supercontinent Gondwana, which was undergoing a long process of separation during the Cretaceous (Ali and Aitchison, 2008; Upchurch, 2008). Therefore, our results speak for a Gondwanan origination of Siphonaptera, as proposed previously by other researchers (Medvedev, 2000a, b). Australia and South America remained terrestrially connected via Antarctica until the late Eocene according to recent estimations (Scher and Martin, 2006; Stickley et al., 2004; Upchurch, 2008; White et al., 2013), permitting an exchange of flea fauna between the two emerging continents. This hypothesis is also supported by divergence time estimates in lineage 2 (Fig. 2), in which the Australian subfamily Stephanocircinae and the South American subfamily Craneopsyllinae split around the Paleocene/Eocene boundary. This scenario was also suggested by previous researchers, although it remained untested until now (Krasnov, 2008; Traub and Starcke, 1980; Whiting et al., 2008).

At the split of lineage 2 (Stephanocircidae) from the rest of the stem Siphonaptera, the likelihood of South America (or the Neotropic) as the ancestral geographic state becomes significantly higher (Fig. 2, Table S4), and continues to be the most significantly supported state until the split of lineages 6 and 7, shortly before the K-Pg extinction. Meanwhile, the likelihood of North America/Nearctic increases with the split of lineage 3 from more derived lineages (Fig. 2, Table S4). This may indicate a series of potential host (and flea) migrations from South America to North America. Geological and paleobiological evidence suggests that these two continents probably had temporary land connections at the Caribbean region during the Late Cretaceous (Beck, 2008; Pascual, 2006), making this migration possible.

The reconstruction result also reveals that fleas expanded to the Old World (Eurasia and Africa) as several independent lineages in the tree. This is reflected in the ancestral state reconstruction of lineage 7, a globally distributed infraorder of extant Siphonaptera. Lineage 3 (Doratopsyllinae, Chimaeropsyllidae and Pulicidae) and part of lineage 5 (Hystrichopsyllinae, Coptopsyllidae, Stenoponiinae and Rhadinopsyllinae) also appear to have become globally distributed since their basal beginnings in South America (Neotropic). These clades include several speciose extant families (such as Ceratophyllidae, Leptopsyllidae and Pulicidae), which may implicate rapid radiations into the new ecological niche (e.g pre-existing hosts) as fleas entered the Old World (Eurasia and Africa).

The route of siphonapteran migration into the Old World deserves further discussion. Given the appearance of North America (Nearctic) at the major nodes, it is possible that fleas traveled from North America to Asia through Beringia or to Europe through Greenland. This route of faunal interchange has also been suggested for terrestrial mammals, which took place around the Paleocene-Eocene boundary (56 Mya) (Beard, 2008). Moreover, the cross-continent similarity of flea faunas in Eurasia and North America has been noticed by flea researchers (Krasnov, 2008; Medvedev, 1996), lending support to this hypothesis.

Another candidate route of dispersal to Eurasia is via Africa, as this state is included (with moderate likelihood) in the ancestral range at the base of lineage 3 (containing Doratopsyllinae, Chimaeropsyllidae and Pulicidae). This hypothesis recalls the generally accepted notion that Pulicidae originated in Africa (Medvedev, 1998; Traub, 1985). Conventional views suggest that Africa either separated from the rest of Gondwanaland as early as Middle Jurassic, or remained connected with South America and East Gondwana until around 80 Mya (Upchurch, 2008). Paleobiological studies provide indications that throughout the period of its isolation, Africa might have had temporary land connections to other Gondwanan continents as well as Laurasia that enabled intercontinental migration of animal fauna (Gheerbrant and Rage, 2006; Upchurch, 2008). Taken together, current knowledge of tectonics does not preclude the plausibility of this “out-of-Africa” route.

Meanwhile, the possibility that the drifting Indian subcontinent carried flea fauna from Gondwana to Asia (consistent with the “out-of-India” hypothesis, see (Ali and Aitchison (2008); Datta-Roy and Praveen Karanth, 2009)) is not supported in this study, as Indomalaya was not recovered to have a high likelihood as an ancestral state at any nodes. Moreover, flea taxa that are endemic to Madagascar (such as *Paractenopsyllus* spp. and *Synopsyllus* spp.), the island thought to be separated from India on its way toward north (Ali and Aitchison, 2008; Upchurch, 2008), do not seem to have close phylogenetic connections with any Indomalayan taxa in our analysis. This suggests that South Asia is probably more of a terminus, rather than a transit station, of flea dispersal.

Combining host and geographic information, the identity of the primary hosts of stem fleas becomes clearer. According to fossil records, Metatheria were present in Laurasia during the Late Cretaceous (Luo, 2003). Recent studies suggest that the ancestors of extant marsupial lineages migrated from North America to South America, and then to Australia during the Late Cretaceous or early Paleocene (Beck, 2008; Luo et al., 2011; Meredith et al., 2008; Nilsson et al., 2010). Therefore, it is possible that stem fleas established an ectoparasitic relationship with early marsupials in Gondwana during the Late Cretaceous, before migrating mostly in the opposite direction, as supported by ancestral state reconstructions in this analysis.

The origin and migration of placental mammals are still controversial topics (Goswami, 2012). While conventional views suggested that Eutheria might have originated in the Southern Hemisphere (Murphy et al., 2001), recent analyses point toward a Laurasian origin of this clade, and a subsequent migration from Laurasia to South America (Hallström and Janke, 2010; O’Leary et al., 2013; Wible et al., 2007). In this case, placental mammals would not have reached the already-isolated Australia until ca. 5 Mya (except for bats, a secondary host group of fleas) (Egerton and Lochman, 2009). Under this scenario it would be unlikely that early fleas had placental mammals as hosts in Gondwanaland during the Late Cretaceous. Therefore, the high likelihood of Eutheria at the base of the Siphonaptera clade as derived from our analysis (see 3.3) might be overestimated due to the more recent radiation of placental mammals and the successful adaptation of fleas (including early lineages) to these new hosts.

### 3.5. Extant fleas, fossil “fleas” and dinosaurs

In the present study the phylogeny, divergence times, ancestral host associations and geographical distributions of extant Siphonaptera were reconstructed, from which the putative historical scenarios of this ectoparasitic insect order can be inferred. Using calibration points within extant fleas, the common ancestor (stem lineage) of extant Siphonaptera began to diverge in the Cretaceous in Gondwanaland. By the Late Cretaceous, fleas were already adapted to an ectoparasitic life style with early Metatheria (early marsupials), and Eutheria, most of which are likely extinct. Early extant flea lineages likely dispersed toward northern continents through a South America-North America-Eurasia route, and perhaps also through a South America-Africa-Eurasia route. Along this path, they encountered and associated with Placentalia, likely subsequent to an earlier start with marsupials. Nevertheless, the main diversification of fleas was associated with the rapid radiation of placental mammals during the Cenozoic, with the most diversified flea families seen in northern continents.

Based on these results, we suggest that that the fossil “flea” taxa discovered from the Mesozoic era of Northeast Asia are unlikely to be phylogenetically clustered within the Siphonaptera clade (Fig. 1), with the following rationale:

First, the earliest divergence (Macropsyllidae) within extant Siphonaptera is averaged into the Late Cretaceous. Given the morphological consistency among all lineages of extant Siphonaptera, it is clear that the unique set of characters used to confer clade affiliation was already present at this time. The youngest (Cretaceous) fossil representatives of both putatively new siphonapteran lineages (Fig. 1, sensu Huang et al., and Gao et al.) overlap not only with the lower tail of the 95% confidence interval (CI) of the estimated onset of extant flea diversification, but more importantly, coincide with the estimated age of origin of Siphonaptera (Early Cretaceous, Table 1A-C). This should lead to caution in interpreting them as part of, or transitioning to Siphonaptera, especially given the distinct morphology of the Mesozoic fossils. Specifically, none of the fossil “fleas” share the following characters with extant fleas: a) a latero-lateral compression of the body, b) winglessness with a modified wing joint (pleural rod, and pleural arch), with enlarged metacoxae and metafemorae, and c) the consistent presence of a saddle-shaped sensilial plate (pygidium), dorsally placed on sternum X (Whiting et al., 2008). In other words, in the context of the estimated Cretaceous origin of extant fleas, there is no clear taxonomic premise to casually include them into what we currently consider Siphonaptera, as has been suggested by Huang et al. (2012, 2013a), and Gao et al. (2012, 2013).

Second, Northeast Asia, where the “flea” fossils were found, represents Laurasia, which was geographically distant from Gondwana, where the stem representative of crown fleas likely originated (also see divergence time estimations). During the Early Cretaceous, Laurasia and Gondwana had already separated from each other by the expanding Tethys Ocean, a geographic barrier that terrestrial animals were unlikely to be able to cross (Hallström and Janke, 2010; Nishihara et al., 2009; Wildman et al., 2007).

Third, therians (marsupials and placental mammals) were recovered to be the most likely hosts of early extant Siphonaptera. Monotremes (which diverged early from Theria) and birds (which are an ingroup of the dinosaur clade), do not have significant representation in the reconstructed ancestral host ranges. Therefore, the scenario that dinosaurs or pterosaurs were primitive hosts of Siphonaptera, as proposed by Huang et al. (2012, 2013a), and Gao et al. (2012, 2013) is not supported.

Finally, although the *Tarwinia* fossil from the Late Cretaceous of Australia (Jell and Duncan, 1986; Riek, 1970) seems to be compatible with the time and location of early modern fleas as recovered in this study, there is no direct evidence to suggest it was already adapted to an ectoparasitic life style, and its evolutionary relationship with modern Siphonaptera is ambiguous (see above).

It is yet too early to draw any definite conclusion regarding the phylogenetic relationship among modern Siphonaptera, *Tarwinia*, Northeast Asia fossil “fleas” and other Mecoptera lineages. However, caution should be taken when describing these Mesozoic giant insect fossils as “pre-fleas” or interpreting them as relatives of, or part of the modern Siphonaptera.

## Acknowledgements

We express our sincere gratitude to the researchers who have contributed specimens to this work as listed in Whiting et al. (2008). In addition, we like to thank Paul Bates, Harrison Institute, Kent, UK; Emmett R. Easton, University of Hawaii at Manoa, Honolulu, HI; Jake Esselstyn, Louisiana State University, Baton Rouge, LA; Alan Harrison, University of Aberdeen, Aberdeen, UK; Si Si Hla Bu and M. Roi Lum, University of Mandalay, Myanmar; John M. Kinsella, Helm West Laboratory, Missoula, MT; Michael Meyers, Christopher Newport University, Newport News, VA; Vasyl V. Tkach, University of North Dakota, Grand Forks, ND; Ulrich E. Schneppat, Bündner Naturmuseum, Switzerland; and Bernard Zonfrillo, University of Glasgow, Scotland. This research was funded by NSF DEB 1050793 awarded to KD, and NSF Career Grant DEB 9983195, awarded to MFW.

## Supplementary information

Tables S1-S4, Figures S1-S2

